# Pigeons exhibit low susceptibility and poor transmission capacity for H5N1 clade 2.3.4.4b high pathogenicity avian influenza virus

**DOI:** 10.1101/2025.05.02.651910

**Authors:** Cecilia Di Genova, Caroline J Warren, Simon Johnson, Sofia Riccio, Kelly Roper, Saumya S. Thomas, Audra-Lynne Schlachter, David Jorge, Kajal Ralh, Jafar Hassan, Elizabeth Billington, Alejandro Núñez, Ian H. Brown, Marek J Slomka, Ashley C. Banyard, Joe James

**Affiliations:** Department of Virology, Animal and Plant Health Agency (APHA-Weybridge), Woodham Lane, Addlestone, Surrey, KT15 3NB, UK; Department of Pathology, Animal and Plant Health Agency (APHA-Weybridge), Woodham Lane, Addlestone, Surrey, KT15 3NB, UK; WOAH/FAO International Reference Laboratory for Avian Influenza, Animal and Plant Health Agency (APHA-Weybridge), Woodham Lane, Addlestone, Surrey, KT15 3NB, UK

**Keywords:** Pigeon, high pathogenicity avian influenza virus, H5N1, HPAIV, transmission, pathogenicity

## Abstract

The ongoing panzootic of H5N1 high pathogenicity avian influenza virus (HPAIV) has caused the deaths of over half a billion wild birds and poultry, and has led to spillover events in both wild and domestic mammals, alongside sporadic human infections. A key driver of this panzootic is the apparent high viral fitness across diverse avian species, which facilitates an increased interface between wild and domestic species. *Columbiformes* (pigeons and doves) are commonly found on poultry premises, yet little is known about their potential role in contemporary HPAIV disease ecology. Here we investigated the epidemiological role of pigeons (*Columba livia*) by determining their susceptibility using decreasing doses of clade 2.3.4.4b H5N1 HPAIV (genotype AB). We investigated infection outcomes and transmission potential between pigeons and to chickens. Following direct inoculation, pigeons did not develop clinical signs, and only those inoculated with the highest dose shed viral RNA or seroconverted to H5N1-AB, revealing a MID_50_ of 10^5^ EID_50_. Even in the high-dose group, only low-level shedding and environmental contamination was observed, and low-level viral RNA were present in the tissues of directly inoculated pigeons, with no distinct pathological lesions. Pigeons did not transmit the virus to pigeons or chickens placed in direct contact. We observed distinct differences in sialic acid receptor distribution in the pigeon respiratory tract compared to chickens and ducks. Together, these findings suggest that pigeons have low susceptibility to clade 2.3.4.4b H5N1 HPAIV and are unlikely to contribute significantly to virus maintenance, transmission to poultry, or zoonotic infection.

## Introduction

Avian influenza viruses (AIVs) are a diverse group of economically important pathogens, classified according to their subtype determined by their haemagglutinin (HA, H1-H16 and H19) and neuraminidase (NA, N1-N9) genes [1, 2]. High pathogenicity AIV (HPAIV) outbreaks in *gallinaceous* poultry (e.g. chickens and turkeys), are typified by rapid onset of high morbidity and mortality [3]. Among HPAIVs, H5Nx viruses of the A/goose/Guangdong/1/1996 (gs/Gd) lineage remain a major global concern and continue to evolve into new clades based on the phylogeny of the HA glycoprotein [4]. In addition, gs/Gd lineage H5Nx viruses have undergone extensive reassortment among the NA and internal genetic segments creating a range of different subtypes and genotypes within each subtype [5]. Europe has experienced clade 2.3.4.4 epizootics since 2014 [6], where clade 2.3.4.4b has been particularly prominent since autumn 2020 [5, 7, 8], with significant wild bird mortality and damaging economic impacts upon the poultry industry [9]. During autumn 2021, H5N1 emerged as the dominant subtype among European clade 2.3.4.4b HPAIVs and has been maintained and disseminated in wild bird populations, particularly aquatic bird species, facilitating spread to the Americas (North America and then South America) via bird migration [10], and onward towards the Antarctic region [11]. This is the largest recorded HPAIV panzootic in terms of detections in wild birds and numbers of poultry outbreaks [12]. Alongside high infection in wild bird and poultry, there have been numerous reports of infections in wild and domestic mammals with clade 2.3.4.4b H5N1, including sustained infection in domestic cattle in the United States of America (USA), and sporadic human infections, which underline the zoonotic potential and pandemic risk [13]. Since 2021, a high degree of genetic diversity has been observed within the H5N1 clade 2.3.4.4b HPAIV, particularly within the internal gene segments though reassortment with local indigenous low pathogenicity avian influenza viruses (LPAIVs). As a result, over 12 different genotypes have been detected in wild birds or poultry in the United Kingdom (UK) alone, including genotype AB among the dominant genotypes, which was also the dominant genotype in continental Europe [5, 14].

A key hallmark of this H5N1 clade 2.3.4.4b HPAIV panzootic is the broad avian host range which the viruses exhibit, and the apparent increased fitness in some wild bird species compared to previous H5Nx gs/Gd HPAIVs [12, 15, 16]. In the UK, there have been extensive infections in seabird populations including *Laridae* (gulls) and *Sulidae* (e.g. gannets) [17, 18]. Among wild bird species, the order *Columbiformes* (pigeon and doves) have been viewed as potential bridging species to captive birds and humans for several infectious diseases [19, 20]. Pigeon behaviour includes foraging on agricultural land, along with colonisation of urban habitats shared with waterfowl (in municipal parks) and humans [21–23]. *Columbiformes* exist as a single family called *Columbidae*, consisting of 41 groups of pigeons and doves, and representing over 300 species [24], and the terms pigeon and dove are used interchangeably for any of the species. *Columbiformes* are very common worldwide, and five species of *Columbidae* are regularly seen in the UK, including the feral pigeon (*Columba livia*), stock dove (*C. oenas*), wood pigeon (*C. palumbus*), collared dove (*Streptopelia decaocto*) and turtle dove (*S. turtur*) [24]. Feral pigeons, common pigeons, racing pigeons, city pigeons and rock doves are all the same species (*C. livia*) which descended from wild rock doves following domestication at least 2000–5000 years ago; they are abundant worldwide, and have high connectivity with human settlements [24].

Historically, *Columbiformes* have been regarded as a relatively unimportant hosts for AIV infections, including HPAIVs [25–27]. However, multiple different subtypes of AIV have been identified in pigeons, including H1, H2, H3, H5, H6, H7, H9 and H11, suggested at least a limited capacity for pigeons to harbour certain influenza viruses [28]. In addition, the emergence and global spread of clade 2.3.4.4b H5N1 HPAIVs since autumn of 2021 has been attributed in part to the broad species range of these viruses, yet relatively little is known about the role of pigeons during this panzootic. Through recent passive wild bird surveillance programmes (targeting wild birds which are found dead) in the UK and continental Europe, few HPAIV H5Nx positive pigeons have been reported [12, 29], suggesting an absence of high mortality in pigeons, but also reducing surveillance samples, and still leaving questions on their infection status. Similarly, infrequent mortalities have been observed in free-living *Columbiformes* in other regions, such as in the USA [30]. However, clade 2.3.4.4b detections appear to be an overrepresented clade in pigeons compared to other H5 clades or subtypes [28]. Therefore, given these considerations, it remains unclear whether subclinical infection and transmission of these HPAIV can occur in pigeons. Their frequent presence at poultry facilities and close association with humans underscore the need to assess the potential role pigeons may play, not only in terms of their susceptibility to contemporary clade 2.3.4.4b HPAIV, but also in determining whether they are capable of sustaining inapparent infections within their populations. This is essential to evaluate their capacity to act as a bridging host for virus transmission to commercial poultry species, such as chickens, and to assess any associated zoonotic risk [19].

Here we selected one of the major European H5N1 clade 2.3.4.4b H5N1 HPAIV genotypes, namely AB (AIV09), which became epidemiologically prominent during 2022 [5, 8]. We used this virus to quantify the susceptibility of pigeons (*C. livia*) to H5N1 clade 2.3.4.4b HPAIV, and to investigate the impact in terms of clinical disease, pathology, tissue tropism, seroconversion and viral shedding dynamics following infection. We also explored the ability of pigeons to maintain infection by assessing pigeon-to-pigeon transmission and assessed the ability of pigeons to act as bridging species to poultry by assessing pigeon-to-chicken transmission.

## Methods

### Virus

The genotype of H5N1 clade 2.3.4.4b HPAIV used for this study was A/chicken/England/014330/2022 (H5N1) (GISAID accession number: EPI_ISL_13370704), herewith referred to as H5N1-AB. This isolate is representative of the dominant genotype which circulated in the UK and continental Europe during the 2021-2023 epizootic and is classified as genotype AB or AIV-09 as defined by the European Union Reference Laboratory (EURL) [31] and Byrne, James [5] respectively. H5N1-AB was propagated in 9-day-old specified pathogen free (SPF) embryonated fowls’ eggs (EFE) and titrated in EFEs to determine 50% egg infectious dose (EID_50_), as previously described [15]. The H5N1-AB stock was diluted in 0.1M pH 7.2 phosphate buffered saline (PBS) to prepare the desired dose for all *in vivo* infections. The precise dose was confirmed by titration of the inoculum in EFEs.

### Animals and pre-sampling

Forty-eight domestic pigeons (*C. livia*) and twenty-four high-health-status Hy-Line Brown chickens (*Gallus gallus*) were sourced from a commercial UK supplier, and Hy-Line UK Ltd (Studley, UK), respectively. The pigeons were quarantined for five weeks in a separate enclosure by the supplier.

Pigeons were approximately three months old at the time of procurement and quarantined for a further two weeks upon arrival in the high containment animal facility. The high-health-status chickens were reared from one-day-old to three-weeks of age in clean environment. Both species were acclimatised for a minimum of seven days prior to infection. The birds were housed under controlled conditions, with temperatures maintained at 21–22°C and relative humidity between 50–60%. Prior to inoculation, approximately 1 ml of blood was collected from the brachial vein of each bird, and oropharyngeal (Op) and cloacal (C) swabs were obtained. All birds tested negative for antibodies against H5N1-AB via haemagglutination inhibition (HI) assay and were also negative for influenza A viral RNA (vRNA) in swab samples.

### *In vivo* study design

To assess a response to three different H5N1-AB inoculation doses, three separate identical rooms were used, with pigeons and chickens randomly allocated into experimental groups. For a single given dose (and room), three groups were used, consisting of; (**i**) directly inoculated pigeons (n=8), (**ii**) contact pigeons (n=8), and (**iii**) contact chickens (n=8) (Figure S1), where the ‘donor’ (D0) and ‘recipient’ (R1) birds were designated as directly inoculated and contact birds, respectively. For each dose (and room), pigeons were housed in “walk-in chicken runs” (Omlet, UK), consisting of a plastic-coated metal cage measuring 2.2 × 2.2 × 1.5 metres. These enclosures were equipped with wall-hanging and floor-located feeding troughs and drinkers, side-mounted apex, and flat perches and nesting boxes. Chickens were housed on the floor of the same cages on autoclaved straw litter, with access to floor-located feeding troughs, drinkers, scratch mats, and boxed shelters. On the inoculation day, the D0 pigeons were separated for six hours from the R1 birds (pigeons and chickens) and inoculated with 0.1 mL of phosphate-buffered saline (PBS) containing H5N1-AB, via the intra-nasal and intra-ocular routes. The majority of the inoculum was administered intra-nasally, and a single drop (∼20 µl) was administered intra-ocularly. The inoculum doses were 10^2^ (low), 10^4^ (medium), and 10^6^ (high) EID_50_. At 3 days post-infection (dpi), two D0 pigeons from each dose group were culled for post-mortem (PM) analysis. All surviving birds were culled at 14 dpi, with PM analysis also performed at this time on three D0 pigeons from the high-dose room only.

### Clinical samples and processing

Op and C swabs were taken daily from all birds from 1 to 14 dpi. Swabs were immersed in 1 mL Leibovitz L-15 Medium (L-15) (Gibco, USA) and stored at −80°C until required for RNA extraction. Blood was collected at 14 dpi by cardiac puncture, under terminal anaesthesia, from all surviving birds. Sera were separated from clotted blood by centrifugation, heat-inactivated at 56°C for 30 minutes and used for downstream serological analyses. At PM a range of tissues were collected including feathers, cervical trachea, pectoralis muscle, heart, liver, pancreas, spleen, kidney, lung, caecal tonsil, bursa, jejunum, caecum, ileum, colon, proventriculus and brain. Tissues were stored in bijoux tubes at −80°C, and a section of each was transferred into L-15 medium and roughly chopped; supernatants were used for RNA extraction. In addition, environmental samples were collected before inoculation and daily from 1-10 dpi, then at 12 and 14 dpi from the high dose room only as drinking water (wall-hanging and floor located drinkers from pigeons and chickens, respectively) and faeces (pigeon faeces were collected from high perches and/or nesting boxes). Additionally, the chicken scratch-mat was swabbed, and a sample of soiled straw-litter taken. Pigeon feathers were also collected daily in the housing environment, with these being identified by their appearance and colour, to determine the species of origin. RNA was extracted directly from all liquid samples, swabs (scratch mat) were processed as described above, and solid samples (faeces, straw litter and environmental-origin feathers) were prepared as a 10% (volume/volume) suspension in PBS (pH 7.2) as previously described [32].

### Clinical scoring and monitoring

Clinical inspections were carried out twice daily from 1 to 3 dpi, then daily until end of study (14 dpi). Birds were monitored against pre-defined clinical score sheets as previously described [15] by experienced animal technicians.

### RNA Extraction and real-time reverse transcription-PCR (RT-PCR)

vRNA was extracted from swabs, environmental samples and PM tissue homogenates, using MagMAX CORE extraction chemistry (Applied Biosystems, Warrington, UK) upon a Kingfisher (Thermo Fisher Scientific, Glasgow, UK) as previously described [15]. For H5N1-AB vRNA detection, matrix (M)-gene RT-PCR primers and probes were used [33]. A ten-fold dilution series of H5N1-AB vRNA at a known EID_50_ was used to establish standard curves using MXPro software (Aria) to determine relative equivalent units (REU), as previously described [15]. For M-gene RT-PCR, Cq values <36.00 (>10^1.98^ REU) were considered as AIV positive, sub-threshold values in the range Cq 36.01–39.99 and Cq 40.00 (No Cq) were discussed, but were interpreted as negative as previously defined [15]. For sample producing a subthreshold Cq value by M-gene RT-PCR, an H5 RT-PCR, which specifically detects clade 2.3.4.4b H5 HPAIV RNA [34] was also performed. An individual bird was considered as productively infected based on at least one positive swab and/or positive seroconversion to H5N1-AB (described below).

### Serology

Sera were tested for H5N1-AB HA reactive antibodies by the haemagglutination inhibition (HI) assay using four haemagglutination units of H5N1-AB. Pigeon sera were pre-absorbed with packed chicken red blood cells prior to testing as previously described [35]. Sera with a reciprocal HI titre ≥16 were considered positive, values <16 are discussed but considered negative according to international standards [35].

### Pathology and immunohistochemistry (IHC)

Following post-mortem, harvested tissue samples were fixed in 10% (v/v) buffered formalin for a minimum period of five days and routinely processed for histopathology, as previously described [36]. Briefly, four-micron thick serial tissue sections were either immunolabelled against influenza A virus NP (Statens Serum Institute, Copenhagen, Denmark), or were stained with haematoxylin and eosin (H&E) as previously described [15]

### Sialic acid receptor distribution analysis in pigeon, chicken and duck tissues

To assess the presence and abundance of α2-3-linked sialic acid (2-3Sia) and α2-6-linked sialic acid (2-6Sia) in respiratory tissues of different bird species, formalin-fixed, paraffin-embedded (FFPE) tissues from three uninfected, healthy individuals per species were examined. Pigeon tissues were collected during the present study. As comparator species, FFPE tissues from chickens (Hy-Line UK Ltd) and Pekin ducks (Cherry Valley hybrid) previously prepared in an earlier study [15] were used. Sections of 4 μm thickness were dewaxed through xylene then rehydrated through absolute alcohol, quenched for endogenous peroxidase using 3% hydrogen peroxide in methanol (VWR International) for 15 min. Slides were assembled into Shandon cover plates to facilitate IHC using the Sequenza system (Thermo Fischer Scientific) and samples underwent a biotin block using Avidin and Biotin solutions (abcam) for 10 mins, followed by a protein block using a 1% BSA (Sigma Aldrich) solution for 20 min. Samples were then incubated for 1 hour with either *Sambucus Nigra* Lectin (SNA) (Vector Labs, B-1305) 1/1500 1.33ug/mL to demonstrate a2-6Sia, *Maakia Amurensis* Lectin I (MAL I) (Vector Labs, B-1315) 1/1500 1.33ug/mL, or *Maakia Amurensis* Lectin II (MAL II) (Vector Labs, B-1265-1) 1/750 1.33ug/mL to demonstrate a2-3Sia-N-acetylneuraminic acid (NAc) and 2-3Sia respectively. This was followed by incubation with ABC (Vector Labs) for 30 mins at and visualised using 3,3-diaminobenzidine tetrahydrochloride (Sigma Aldrich) for 10 min. “Lectin TBST” (20mM Tris Base, 100mM NaCl, 1mM CaCl_2_, 1mM MgCl_2_ in dH_2_O, adjusted to pH 7.2 using HCl) was used for rinsing sections between incubations and to dilute the biotinylated lectins, ABC and block. Sections were then counterstained within Mayer’s haematoxylin (Pioneer Research Chemicals Ltd.), dehydrated and cleared in absolute alcohol and xylene, and glass coverslips mounted using ClearVue Mounting Medium (Epredia). To confirm the specificity of the lectin binding, matching serial sections were incubated in 100uL of Neuraminidase (Roche) overnight at 37°C following the protein step block and prior to the incubation with the lectins. Presence of histochemical signal in respiratory epithelial cells was evaluated semi-quantitatively on a scale of - (no labelling) to ++++ (abundant labelling).

### Statistical analysis and determining minimum infectious dose (MID)

All statistical analyses were performed using Prism version 8.4 software (GraphPad, San Diego, CA, USA), as indicated on the corresponding figure. The minimum infectious dose (MID) and 50% MID (MID_50_) were calculated as previously described [37].

## Results

### A high dose of H5N1-AB is required to establish infection and seroconversion in pigeons

To assess the virulence, pathogenesis, and transmission characteristics of H5N1-AB in pigeons, groups of eight D0 pigeons were inoculated with one of three H5N1-AB doses; low (10^2^ EID_50_), medium (10^4^ EID_50_), or high (10^6^ EID_50_).

Pigeons inoculated with the low or medium doses did not shed H5N1-AB vRNA from the Op or C cavities (Fig. 1A, B, D & E). A single pigeon in the medium-dose group at 2 dpi exhibited a single subthreshold level of vRNA from in the OP cavity (Fig. 1B), but this was not detectable using an H5 RT-PCR. In contrast, vRNA was shed by seven of the eight pigeons (87.5%) in the high-dose group between 1 and 7 dpi (Fig. 1C, F & G). However, OP shedding levels remained low, with mean peak vRNA titres of 10^3.74^ REU, and an individual maximum titre reaching 10^4.12^ REU. Cloacal shedding was comparatively lower and more sporadic, only a single C swab was positive for vRNA from pigeon #48 at 5dpi with a titre of 10^2.41^ REU. Four of the eight pigeons exhibited subthreshold cloacal shedding, all these were also detectable by the H5 RT-PCR (Fig. 1F & G). No clinical signs were observed in the D0 pigeons throughout the duration of the study (1-14dpi), except for one pigeon in the high-dose group (Pigeon #43) at 1 dpi. This pigeon displayed transient clinical signs which included shivering and a hunched posture, with lowered tail and wings (Table S1).

**Fig. 1.**
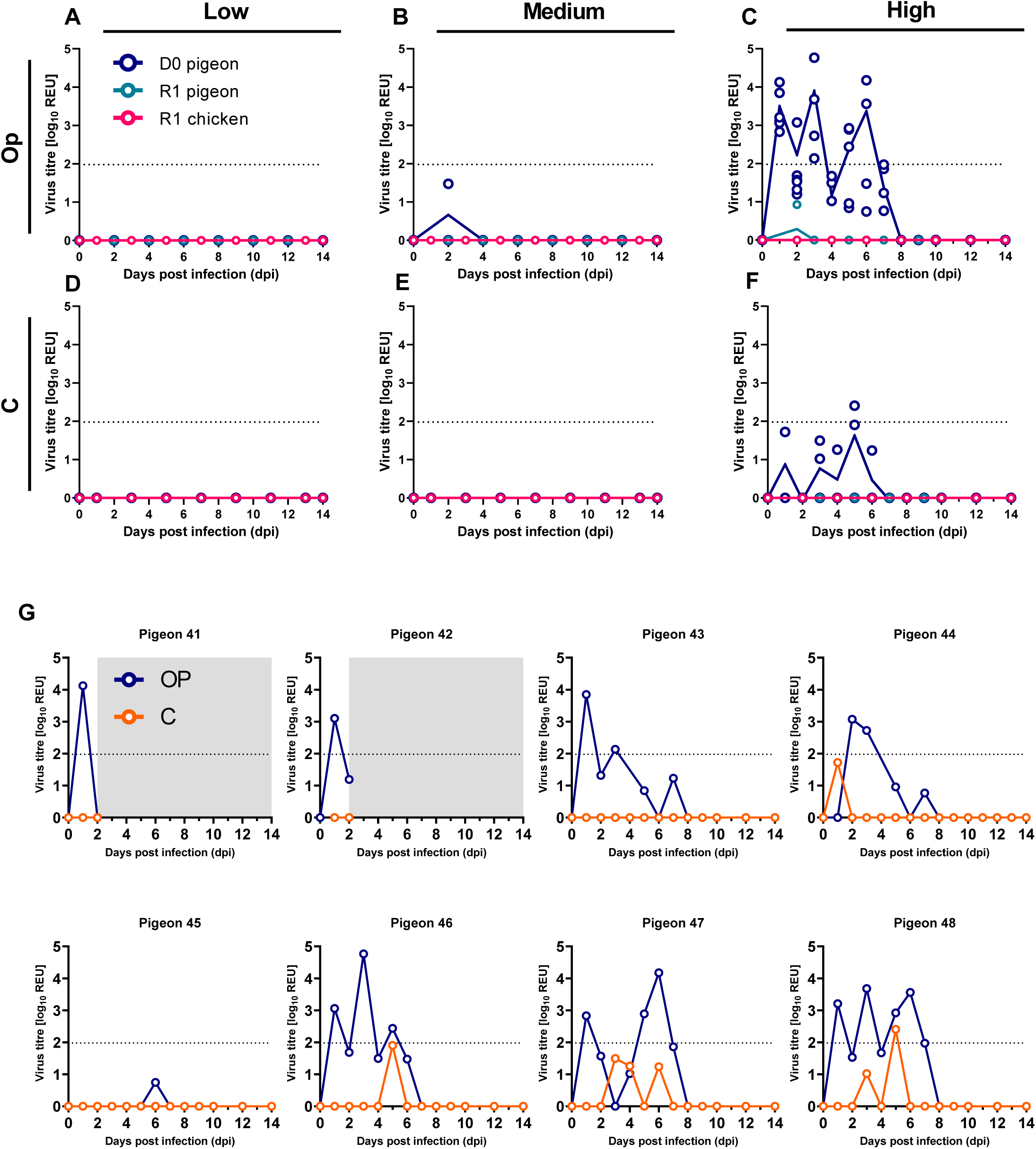
H5N1-AB vRNA shedding from directly inoculated pigeons and contact pigeons and chickens. **A-F.** H5N1-AB vRNA shedding from pigeons which were directly inoculated (D0, blue) with low (10^2^ EID_50_) (**A & D**), medium (10^4^ EID_50_) (**B & E**) or high (10^6^ EID_50_) (**C & F**) doses of H5N1-AB. Shedding is also shown from pigeons (teal) and chickens (pink) placed in direct contact (R1) with D0 pigeons. Shedding is shown from swab samples collected from the oropharyngeal (Op) (**A-C**) and cloacal (C) (**D-F**) cavities. **G.** H5N1-AB shedding from each individual D0 pigeon infected with the high dose (10^6^ EID_50_) is presented as separate graphs for Op (blue) and C (orange). Grey shading indicates no data generated as these pigeons were culled for post-mortem analysis at 3dpi. **A-G**. Viral RNA titres were determined by M-gene RT-PCR and converted into relative equivalency units (REU). Dotted horizontal lines indicate the positive threshold at 10^1.98^ REU.

Consistent with the absence of vRNA shedding, pigeons in the low- and medium-dose groups did not seroconvert, as evidenced by HI titres <1/16 against homologous H5N1-AB at 14 dpi (Fig. 2A & B). In contrast, all pigeons in the high-dose group seroconverted, demonstrating detectable antibodies to H5N1-AB by HI (Fig. 2C & D).

**Fig. 2.**
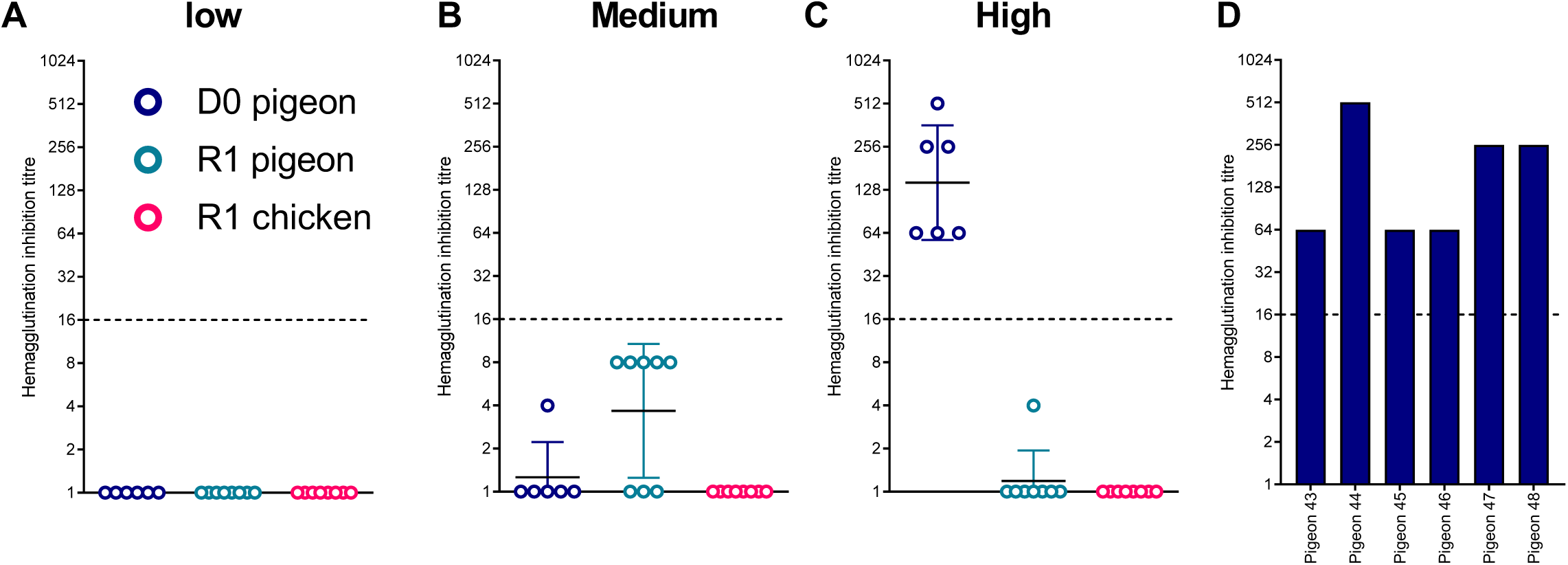
Seroconversion of pigeons directly inoculated with H5N1-AB and from pigeons and chickens placed in direct contact. Haemagglutinin inhibition (HI) titres from sera collected at 14 days post infection (dpi) from pigeons which were directly inoculated (D0, blue) with low (10^2^ EID_50_) (**A**), medium (10^4^ EID_50_) (**B**) or high (10^6^ EID_50_) (**C**) doses of H5N1-AB, or contact (R1) pigeons (teal) and chickens (pink) cohoused with each group. HI titres were derived using homologous H5N1 HPAIV antigen. Graphs show geometric mean **±** geometric SD. **D.** Individual HI titres plotted for pigeons directly infected with the high (10^6^ EID_50_) H5N1-AB. Dotted horizontal lines indicate the positive threshold at 10^1.98^ REU.

Together from shedding and seroconversion data for D0 pigeons, a dose of 10^2^ or 10^4^ EID_50_ resulted in no detectable infection in pigeon (n=0/6), whereas a dose of 10^6^ EID_50_ resulted in 100% infection (n=6/6). The estimated MID_50_ for H5N1-AB in pigeons was therefore >10^4^ but ≤10^6^ EID_50_, and the calculated MID_50_ was 10^5^ EID_50_.

### Transmission of H5N1-AB is ineffective between pigeons and from pigeons to chickens

To investigate the transmission efficiency and dynamics of H5N1-AB within species (pigeon-to-pigeon) and between species (pigeon-to-chicken), we introduced, and cohoused, contact (R1) pigeons and chickens to each dose group (low, medium and high).

Pigeon-to-pigeon transmission efficiency was determined to be 0% (0/8) across all three dose groups based on H5N1-AB vRNA shedding from the R1 birds (Fig. 1A–F, teal symbols). One R1 contact pigeon (pigeon #39) in the high-dose group exhibited a single subthreshold level of vRNA in an Op sample at 2 dpi (10^0.929^ REU) (Fig. 1C), although no vRNA was detected in this sample using the H5 RT-PCR. Additionally, none of the contact pigeons in any dose group seroconverted to H5N1-AB (Fig. 2A–C). Among the R1 pigeons, negative but subthreshold HI titres (<16 HI) were detected in five of eight (62.5%) and one of eight (12.5%) pigeons cohoused in the medium- and high-dose groups, respectively (Fig. 2B & C). None of the R1 pigeons across all dose groups exhibited clinical signs until study-end at 14 dpi with the exception of one R1 pigeon (#01) in the low-dose group (which exhibited huddling and altered body position) (Table S1).

Pigeon-to-chicken transmission efficiency was similarly 0% (0/8). None of the contact R1 chickens in any dose group shed detectable H5N1-AB (Fig. 1A–F, pink symbols), nor did they seroconvert to H5N1-AB (Fig. 2B–C). Additionally, no clinical signs were observed among any of the R1 chickens during the study period (up to 14 dpi) (Table S1).

### Limited environmental H5N1-AB contamination resulted from pigeon infection

We also investigated the extent of environmental H5N1-AB contamination in the high-dose group by collecting various samples throughout the study period (1–14 dpi). Only a single sample from floor-level drinking water at 5 dpi was positive for vRNA, albeit at low levels with a titre of 10^2.32^ REU. All other environmental samples were negative (either undetectable or subthreshold). Subthreshold vRNA titres were obtained from a single pigeon faecal sample collected at 3 dpi, and a sample collected from the floor-level scratch mat at 7 dpi (Fig. 3). Both these subthreshold detections of vRNA were also detectable by the H5 RT-PCR. No vRNA was detected in the elevated drinking water, pigeon feathers, bedding, or chicken faeces.

**Fig. 3.**
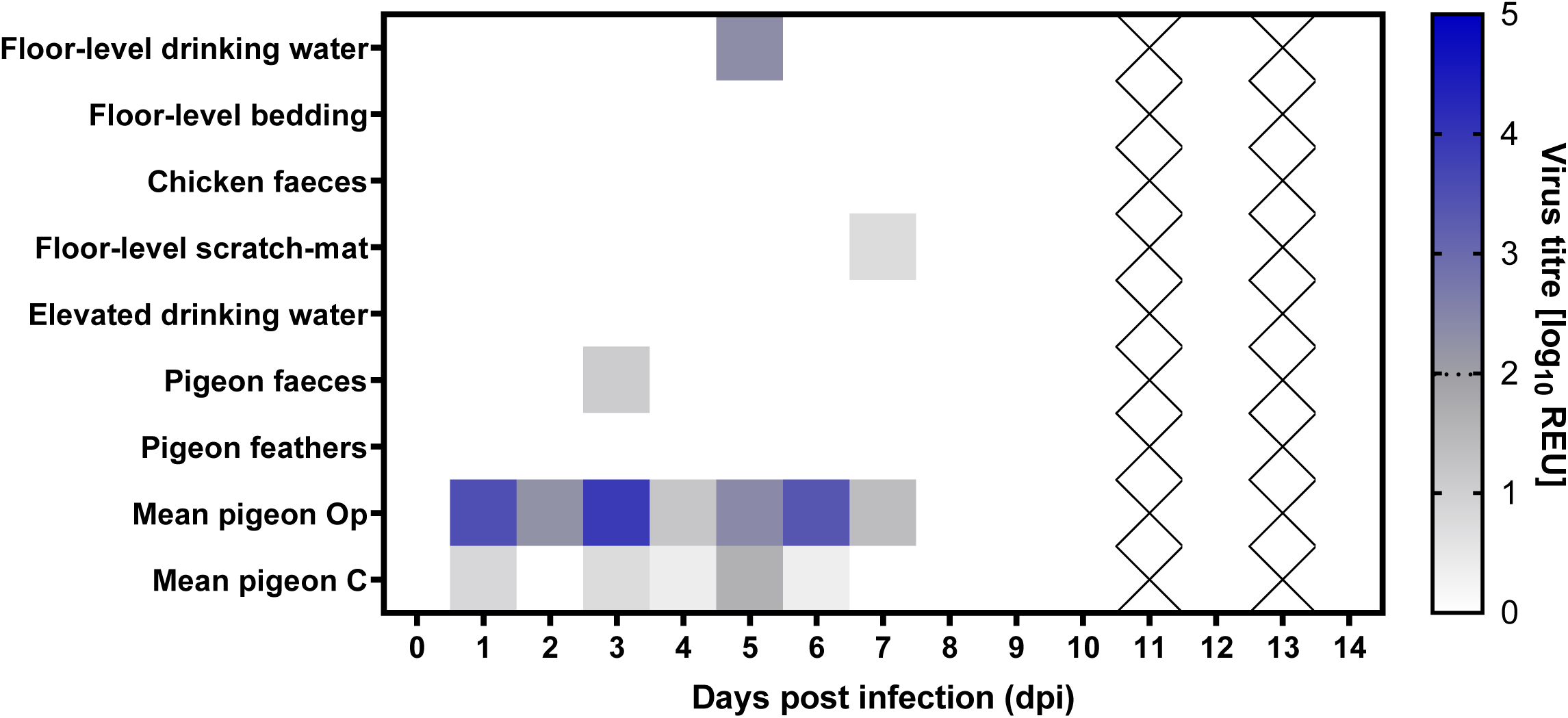
Detection of H5N1-AB vRNA in the environment of pigeons and chickens. Indicated samples were collected from the environment of pigeons directly inoculated with a high dose (10^6^ EID_50_) of H5N1-AB, which also contained direct contact pigeons and chickens. Environmental samples associated with either chickens or pigeons were collected on indicated days post infection (dpi). H5N1-AB vRNA titres were determined by M-gene RT-PCR, with Cq values converted into relative equivalency units (REUs) and displayed graphically as a heat map with high values represented with more intense colouring. Values below the threshold are shown in shades of grey, values above the positive threshold are shown in shades of blue; dotted line in the heatmap indicates the positive threshold at 10^1.98^ REU. Cells with crosses indicate no sample obtained. Mean Op and C shedding from the D0 pigeons is also indicated.

### H5N1-AB did not exhibit widespread tissue dissemination in infected pigeons

The tissue tropism of the H5N1-AB was evaluated in pigeons in the high dose group following direct inoculation. vRNA levels were quantified in tissue extracts obtained from various organs of two D0 pigeons at 3 dpi (pigeons #41 and #42) and three D0 pigeons at 14 dpi (pigeons #43, #46, and #47) (Fig. 4). At 3 dpi, vRNA was only detected in tissues from pigeon #41 while pigeon #42 was negative across all tissues. In pigeon #41, vRNA was detected in the brain, caecal tonsil, caecum, colon, heart, kidney, lung and proventriculus; the highest vRNA levels were found in the brain, caecal tonsil, and lung, with titres of 10^4.02^, 10^3.33^, and 10^3.32^ REU, respectively (Fig. 4). Subthreshold levels of vRNA were identified in the bursa, cervical trachea, jejunum and spleen, vRNA was also detectable in these samples by the H5 RT-PCR. In contrast, only subthreshold levels of vRNA were identified in the lung and spleen of pigeon #42 (Fig. 4), although these were also detected by an H5 RT-PCR. By 14 dpi, at study-end, vRNA was undetectable in tissues from all three pigeons (pigeons #43, #46, and #47) (Fig. 4).

**Fig. 4.**
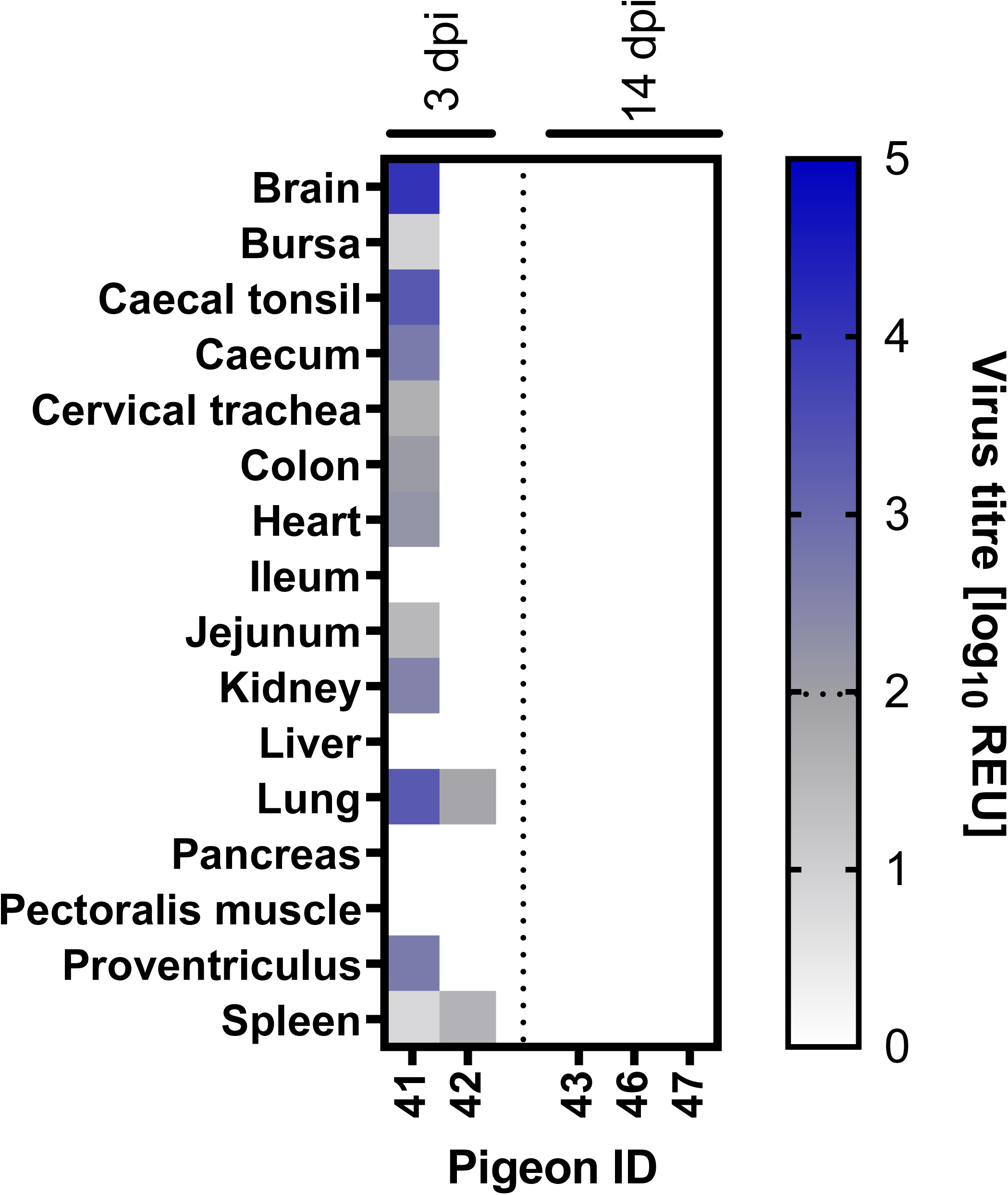
H5N1-AB vRNA distribution in tissues from inoculated pigeons. Tissues were dissected from two pigeons at 3 days post infection (dpi) and three pigeons at 14 dpi following infection with a high (10^6^ EID_50_) dose of H5N1-AB. Viral RNA titres were determined by M-gene RT-PCR, converted into relative equivalent units (REUs) and displayed graphically as a heat map, with higher titres represented by more intense colouring. Subthreshold viral RNA levels are shown in shades of grey, levels above the positive threshold are shown in shades of blue; dotted line in the heatmap indicates the positive threshold at 10^1.98^ REU.

To further characterise tissue tropism, IHC analysis and histopathological examinations were performed on all the tissues obtained from the same five D0 pigeons in the high-dose group, including the two pigeons at 3 dpi (#41 and #42) and three pigeons at 14 dpi (#43, #46, and #47). Histopathological investigations revealed mild multifocal acute airsacculitis in pigeons #41 (3 dpi), #43, and #47 (both 14 dpi). Pigeon #41 additionally exhibited mild multifocal acute dermatitis, moderate multifocal folliculitis of the feather pulp, and moderate multifocal pale irregular areas in the pancreas, suggestive of post-insult regeneration of the parenchyma. No other pathological lesions were identified in any of the examined tissues. IHC analysis did not reveal detectable viral nucleoprotein staining in any of the tissues assessed (Fig. S2).

### Pigeons have lower 2-3Sia and higher 2-6Sia abundance in their respiratory tract compared to chickens and ducks

H5N1-AB has a receptor preference for 2-3Sia compared to 2-6Sia [38], therefore we compared 2-3Sia and 2-6Sia receptor distribution across respiratory tissues (nasal turbinates, trachea and lungs) of pigeons, chickens and ducks and using lectin histochemistry, using the lectins SNA (binds 2-6Sia), and the lectins MAL I and MAL II (binds 2-3Sia) (Fig. S3). In the nasal turbinates and trachea, pigeons showed weaker MAL I and MAL II (2-3Sia) staining, whereas chickens and ducks display much stronger MAL II (2-3Sia) labelling (Table 1). In contrast, pigeons exhibit very strong SNA (2-6Sia) labelling in the nasal cavity, while ducks show no detectable staining (Table 1). In the pulmonary airways, pigeons exhibit strong SNA (2-6Sia) and MAL I (2-3Sia) binding, but ducks and chickens often show even higher or comparable levels of staining, particularly ducks with robust MAL II (2-3Sia) and SNA (2-6Sia) labelling (Table 1). A large difference was seen in the pulmonary air capillaries where pigeons lack MAL I (2-3Sia) staining entirely and show only moderate MAL II (2-3Sia) with strong SNA (2-6Sia) labelling. In contrast, ducks show robust MAL II (2-3Sia) and moderate SNA (2-6Sia) staining, while chickens have weaker overall staining (Table 1). Together, these comparisons suggest that pigeons have a relatively lower presence of 2-3Sia but higher 2-6Sia abundance in upper respiratory tissues compared to chickens and ducks.

**Table 1.**
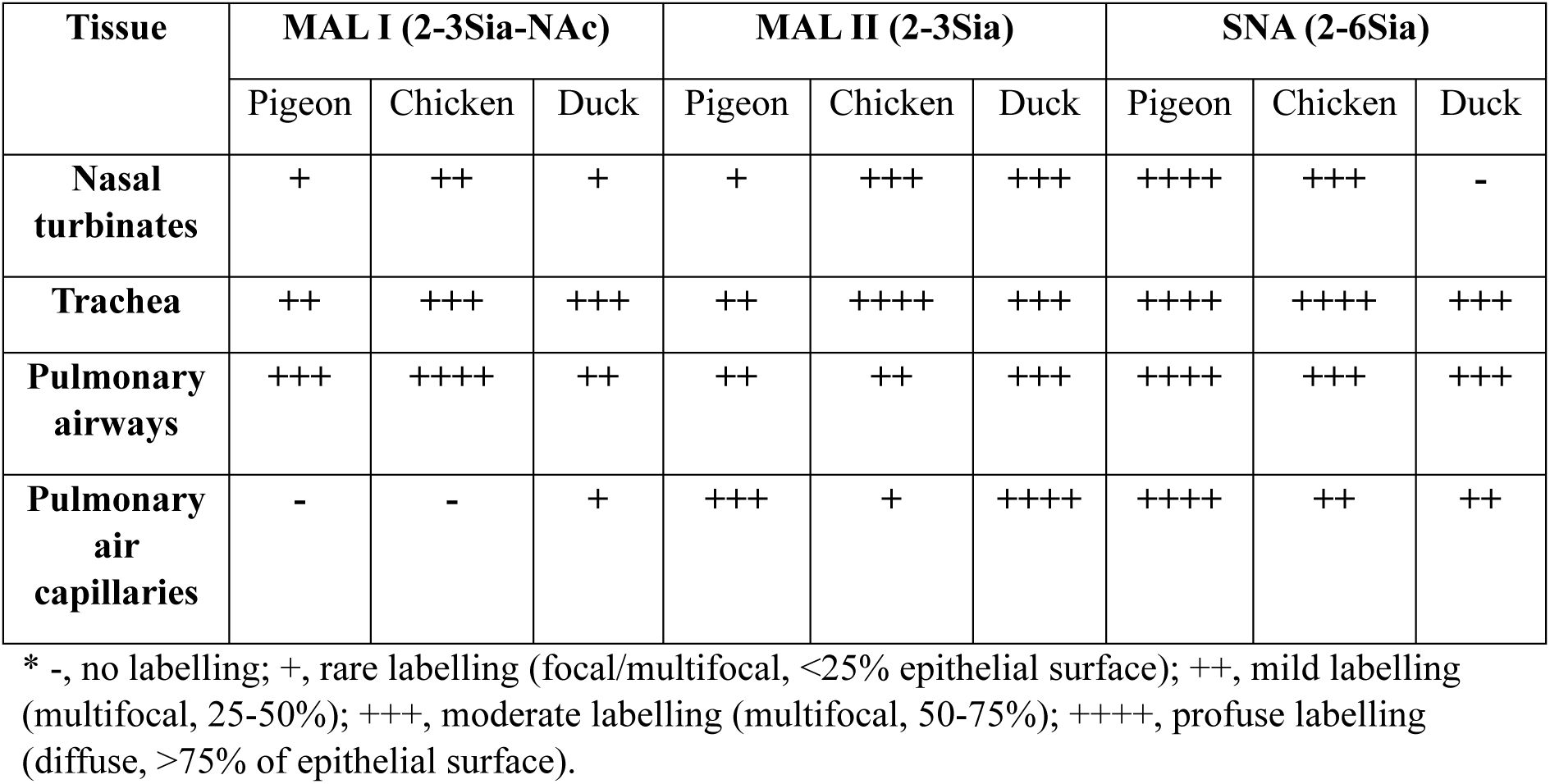
Distribution of sialic acid receptors by lectin histochemistry in respiratory tissues of pigeons, chicken and ducks.

## Discussion

The ongoing panzootic which started in 2021 is the largest H5Nx HPAIV outbreak on record for Europe and the Americas in terms of number of infected wild birds and poultry incursions, with the scale and extent of spread attributed to enhanced fitness in wild birds, and the broader avian species range [15, 39]. In this study we demonstrated the limited susceptibility, absent inter- and intra-species transmission, and low-level tissue dissemination in pigeons (*C. livia*) following inoculation with a contemporary European H5N1 clade 2.3.4.4b HPAIV (genotype AB). This study clearly showed that pigeons are unlikely to act as a direct bridging species due to productive infection for the transmission of this virus to chickens, as an important commercial poultry species. However, their role as an indirect bridging species, through fomite transmission, was not assessed in this work and requires further study. Our findings indicate that a high infectious dose (10^6^ EID50) of H5N1-AB is required to cause detectable infection in the directly inoculated (D0) pigeons. Infection among all eight (100%) high dose D0 pigeons was confirmed by vRNA shedding and/or positive serology. D0 pigeons inoculated with low (10^2^ EID_50_), or medium (10^4^ EID_50_) doses exhibited no detectable vRNA shedding and no seroconversion, demonstrating the limited susceptibility of pigeons to these viral doses. Even when pigeons were infected in the high-dose group, shedding was minimal, was predominantly detected from OP samples, and was short-lived (up to 7 dpi). In some instances, our analyses included subthreshold detections of vRNA by M-gene RT-PCR (vRNA titres >Cq 36 or <10^1.98^ REU). While we have presented these data and attributed REU values, these detections arise late in the RT-PCR amplification. In these instances, we also tested the RNA using an H5 RT-PCR assay which detects vRNA from a different gene segment to further increase our confidence in detection and quantification of vRNA. Indeed, previous H5N1 gs/Gd HPAIV outbreak investigations have shown that such high Cq value samples, as measured by M-gene RT-PCR testing, are unlikely to represent infectious virus [40, 41]. Therefore, these subthreshold detections are of little significance. Indeed, this consideration was reflected in the absence of pigeon-to-pigeon transmission and pigeon-to-chicken transmission with H5N1-AB, which correlated with low viral environmental contamination. Environmental contamination has been consistently shown to be associated with efficient H5Nx clade 2.3.4.4b HPAIV transmission among ducks [15, 37] and pheasants [42], with viral environmental contamination reported during UK H5N1 HPAIV poultry farm outbreaks in recent years [32]. These low levels of shedding, environmental vRNA, and absence of transmission to either pigeons or chickens further underscored the limited potential for pigeons to act as significant amplifiers of H5N1 HPAIV in shared habitats with other avian species.

Several studies have historically conducted experimental infection studies with AIV in pigeons. A review published in 2014 summarised the findings from 22 experimental pigeon infection studies conducted with a range of LPAIVs and HPAIVs and concluded that minimal mortalities and low-level virus shedding over brief periods were generally observed in pigeons [19]. Specifically, investigations with earlier gs/Gd lineage H5Nx HPAIVs have also shown similar results; experimental infection with clade 2.2 and clade 2.3.2.1 H5N1 HPAIVs demonstrated low levels of viral shedding [43]. Studies with H5N2, H5N8 and H5N6 clade 2.3.4.4 HPAIV also found inconsistent, short-term and at low viral shedding viral shedding [44–47]. At most, these studies with earlier H5Nx clade 2.3.4.4 HPAVs reported infrequent pigeon mortalities. Recently, and comparably to our study, Root et al (2024) investigated infection and transmission dynamics in common pigeons (*C. livia*) following experimental infection with a contemporary American clade 2.3.4.4b H5N1 HPAIV, representative of the North American epizootic [48]. Root et al (2024) used a dose of 10^6.6^ plaque forming units (PFU), in excess, but broadly equivalent to the high dose used in this study (10^6^ EID_50_), to inoculate eight pigeons, some of which failed to shed virus and only a single bird shed infectious levels exceeding 10^3^ PFU. Six of eight (75%) inoculated pigeons demonstrated seroconversion [48].

The maximal shedding titres observed by Root et al. (2024) are broadly equivalent to those reported in our study (10^3^ PFU compared to 10^4.12^ REU)[48]. Importantly both of these titres are considerably lower than those shed from the Op cavity of ducks and chickens following experimental infection with a similar clade 2.3.4.4b H5N1 HPAIV from the same epizootic, which typically range from 10^4^ to 10^7^ REU in ducks and 10^4^ to 10^6^ REU in chickens [15]. Consistent with the low levels of vRNA shedding from infected pigeons in our study, we also observed very low levels of vRNA in the environmental sources, particularly in the drinking water; whereby water sources have previously correlated with detectable levels of vRNA and the potential for transmission [15]. In our study, low levels of vRNA were detected only in a single sample of drinking water. Water contamination from ducks infected with a similar European clade 2.3.4.4b H5N1 HPAIV, from the same epizootic, was consistently positive during the shedding period, with maximal titres of 10^4.60^ REU [15]. Therefore, water contamination by pigeons was not only significantly less frequent, but also >2 log_10_ lower (10^2.32^ REU) compared to than from infected ducks. By considering the proportions of D0 pigeons infected by a given dose, the MID_50_ of H5N1-AB was calculated as 10^5^ EID_50_. While there may be some genotype variation, this pigeon MID_50_ is comparatively higher than MID_50_ values calculated from experimental infections of other avian species with similar clade 2.3.4.4b viruses, including ducks (<10^2^ to 10^3^)[15, 37, 49] and chickens (10^3.2^ to 10^5^) [15, 50–52]. Overall, these studies indicate that pigeons are less susceptible to H5 clade 2.3.4.4b HPAIV infection, compared to ducks and chickens.

The low vRNA shedding titres and negligible environmental contamination coupled with the relatively high MID_50_ in pigeons likely explains the lack of transmission we observed between pigeons in this study. Root et al. (2024) also investigated transmission from 8 directly infected pigeons to two contact pigeons, both (100%) of the contact pigeons shed virus on a single day post contact, and seroconverted via HI, suggesting some transmission had occurred. This observed transmission difference from our study may be explained by housing density or experimental design differences, which may have altered the frequency or degree of contact between pigeons. Equally, despite both viruses being clade 2.3.4.4b H5N1 HPAIV representatives of the current panzootic, these viruses differ in their internal genes and so impact of genotype must be considered. We selected an AB genotype representative of the dominant genotype detection in the UK and continental Europe [5, 31], whereas Root et al. (2024) selected an American H5N1 HPAIV genotype. Soon after the start of the European H5N1 HPAIV epizootic in 2021, the virus was translocated into North America via bird migration where it underwent extensive reassortment with indigenous American LPAIVs. Thus, genetic differences within the virus may have different propensities for infection and transmission in pigeons. However, the general observations of a lack of clinical signs and low shedding titres were consistent between the two genotypes in pigeons.

Pigeons are frequently observed both on and near poultry farms and have a high connectivity with human activity; with confirmation that pigeons were one of the most frequently observed avian species on poultry farms using camera trap studies [53]. In addition, air sampling from directly inside the poultry houses affirmed presence of genomic DNA from *Columbiformes* [54], indicating strong interactions between pigeons and poultry. Therefore, we investigated the transmission risk from pigeons to chickens, through co-housing. Consistent with the low virus shedding in pigeons, and lack of environmental contamination and pigeon-to-pigeon transmission, we observed 0% transmission to co-housed chickens, based on shedding and seroconversion in the contact chickens. One previous study with H5N6 clade 2.3.4.4 HPAIV similarly showed 0% transmission from infected pigeons to chickens [46]; however, this has not been assessed by any study with contemporary H5N1 HPAIV, but indicates a limited ability of pigeons to act as bridging species for poultry incursions of H5Nx clade 2.3.4.4b HPAIVs. While in this study we investigated the ability of pigeons to directly transmit H5N1 clade 2.3.4.4b to chickens and pigeons, we did not assess the role pigeons may play as ‘vectors’ in indirect transmission, as this was outside of the scope of this study. Rodents have been hypothesised to act in this way though mechanically transferring contaminated material into poultry houses [55]. It is conceivable that pigeons and *Passeriformes* (perching birds) may also act as vectors for fomite transmission, although this requires further study. However, the limited risk of pigeons to act as bridging species is supported by a lack of association being observed in the UK between *Columbiformes* distribution and infected poultry premises during the 2021-2023 panzootic, when spatial generalized additive models were used based on public bird observations reports [56].

Following direct infection of pigeons, we observed no clinical signs and no mortalities; this finding is broadly consistent with the outcomes of other similar experimental pigeon infection studies [44, 45, 47, 48]. We detected overall low vRNA tissue titres, but viral dissemination occurred in one of the two pigeons at 3dpi, with the highest vRNA titres in the brain, lung, and caecal tonsil. Importantly, no nucleoprotein was detected in any tissue by IHC. We also did not observe any histopathological lesions associated with viral associated tissue damage. Together, these results indicated that while vRNA was present at low levels, H5N1-AB was not replicating to high levels in these organs, suggesting that, although systemic dissemination of the virus can potentially occur in pigeons, it is highly variable between pigeons and does not result in efficient replication producing high virus titres. By 14 dpi, no vRNA was detected in any tissues, suggesting efficient viral clearance in pigeons [57]. A similarly low level of virus dissemination in pigeon tissues has been previously identified following infection with earlier gs/Gd clades [43]. However, there do appear to be virus strain-specific differences in terms of infectivity and infection outcome in pigeons. Lui et al (2022) performed pigeon experimental infection studies with two clade 2.3.4.4 H5N6 HPAIVs [46]. One strain caused no clinical disease with minimal viral shedding, whereas the other strain caused 25% mortality, with 58% of pigeons displaying neurological signs and wide dissemination in tissues with this strain [46]. In addition, there have been several documented reports describing infection and mortalities of *Columbiformes* with H5Nx HPAIV, including common pigeons [58], wood pigeons [59] and African mourning doves (*S. decipiens*) [60]. A recent case of natural HPAI H5N1 clade 2.3.4.4b infection was recorded in a wood pigeon (*C. palumbus*) found deceased in a wildlife centre in Germany [59]. Gross examination revealed mild splenomegaly and discolouration of the pancreas, while histopathological analysis identified neuronal necrosis within the grey matter of the cerebral hemispheres and brainstem, demonstrating pronounced neurotropism of the virus [59]. This wood pigeon mortality may have been due to a greater individual susceptibility to systemic H5N1 HPAIV infection, with resultant death or the ability of different genotypes to infect. Similarly, natural infection of pigeons with an older H5N1 clade 2.2.1 HPAIV in Egypt was associated with 50% mortalities, with congestion of the internal organs, particularly the lungs and brain being observed [58]. Interestingly, when an isolate from this outbreak was assessed in experimental pigeon infections, some clinical signs did ensue, but with reduced mortality (10%) compared to the field observations [58]. Passive surveillance of found dead wild birds in the UK since the start of the clade 2.3.4.4b H5N1 HPAIV panzootic in October 2021 has shown H5 HPAIV infection in only 23 *Columbidae* out of 3465 (0.66%) [16].

These observations suggest that while H5 HPAIV infection can be fatal in pigeons under certain conditions, mortality is more likely to occur in field settings. This may be attributed to differences in exposure dose, route of infection, or repeated exposure, which may be different in natural environments. Additionally, co-morbidities and environmental stressors likely increase pigeon susceptibility to infection and contribute to more severe disease outcomes. Such factors may also account for the variability observed across experimental studies. Since pigeons are not routinely raised under specific pathogen-free (SPF) conditions and are often sourced from natural or commercial populations, their baseline health status and infection history are typically unknown and heterogeneous. Furthermore, due to the wide range of possible subclinical infections and practical limitations, this information is rarely comprehensively reported in the published literature. Indeed, pigeons in the wild have a high propensity for infection with a range of bacterial and viral pathogens [20, 61]. One of the most significant is avian paramyxovirus type 1 (APMV-1), also referred to Newcastle disease virus (NDV) when virulent forms are seen in poultry [62, 63]. APMV-1 have specific lineages referred to as pigeon paramyxovirus type 1 (PPMV-1) in pigeons, and while prevalence is difficult to assess, between 5 - 17% common pigeons have been found as actively infected, dependent on season [64]. The pigeons used in our study were confirmed negative for APMV-1 infection. However, interestingly co-morbidities between HPAIV and PPMV-1 have been hypothesised for exacerbated mortality in natural infected pigeons in South Africa [65], although the interaction between co-infections and impact on disease outcomes require further study.

Despite these sporadic instances of mortality, most surveillance and experimental infection data, including from this study, have shown that pigeons have a low susceptibility to H5Nx clade 2.3.4.4b HPAIVs, and indeed many other AIVs. The underlying reason behind this is likely multi-factorial. For a species to be susceptible to AIV, the target host cell must be ‘susceptible’ (possess the corresponding receptor for the virus) and ‘permissive’ (possess the corresponding host factors to allow viral replication). Influenza viruses bind to sialic acid and the majority of AIV have a preference for 2-3Sia, instead of 2-6Sia [38]. Therefore, to investigate susceptibility, we assessed the 2-3Sia and 2-6Sia receptor abundance on pigeons using chickens and ducks as a comparator susceptible species. We observed a lower relative 2-3Sia, and higher 2-6Sia abundance in proximal respiratory tissues of pigeons when compared with chickens and ducks. This observation aligns with previous analyses of pigeon tissues which demonstrated that the epithelial surfaces of the larynx, trachea, bronchus, and bronchiole of pigeons contained little or no 2-3Sia, but abundant 2-6Sia distribution [66]. In contrast, chicken, turkey and duck respiratory tissues demonstrated a predominance for 2-3Sia [66, 67]. The vast majority of avian origin clade 2.3.4.4b H5N1, including the virus used here (H5N1-AB), have been shown to exclusively bind 2-3Sia in biophysical analysis, with no detectable binding to 2-6Sia [38]. While more detailed investigations of the pigeon glycome are required, these observations offer a potential mechanism for their low susceptibility to H5N1 clade 2.3.4.4b HPAIVs, and potentially many other AIVs.

With respect to the permissiveness of pigeon cells to clade 2.3.4.4b H5N1 HPAIV, the paucity of pigeon-specific reagents hampers detailed investigation of AIV replication dynamics and associated host responses in these cells. Consequently, this aspect remains largely unexplored. However, analysis of the pigeon immune response to HPAIV has found relatively little change in either cytokine production [57] or proinflammatory factors [43] following infection with H5 HPAIV. Interestingly, it has been noted that pigeons appear to have a higher basal expression of interferon stimulated genes (ISG) compared to other avian species [43]. This observation may be linked to different efficiencies of pigeon MDA5 to sense PAMPs, and eliciting an interferon response, compared to other birds [68]. Therefore, it is also possible that higher basal ISG expression may also contribute to the low pigeon susceptibility to infection.

Under the prevailing circumstances of the ongoing H5Nx clade 2.3.4.4b panzootic [69], the negligible role of pigeons in active disease ecology including direct transmission of these viruses is at least reassuring, particularly given their widespread presence in urban environments and agricultural landscapes. While the virus can infect pigeons at high doses, their limited shedding, lack of efficient transmission to co-housed birds, and minimal environmental contamination suggest they are unlikely to contribute significantly directly to the epidemiology of this virus in wild bird populations or commercial poultry. However, the indirect role pigeons may play though fomite transmission (i.e. bringing contaminated material into a poultry premises) requires further investigation. These findings have important implications for surveillance and control strategies, as they suggest that targeted interventions should focus on species with higher susceptibility and transmission potential, such as wild waterfowl, seabirds, and poultry. Further research into other *Columbiformes* species and their interactions with high-risk avian hosts could provide additional insights into their role, if any, in the spread of H5N1 HPAIV.

## Author statements

### Ethics statements

The experiments were approved by the Animal and Plant Health Agency (APHA) Animal Welfare and Ethical Review Body (AWERB), ensuring compliance with UK legislation and alignment with UK Home Office project licence PP7633638 and PP9307748. All work involving infectious HPAIV was performed in licensed Containment Level 3 (CL-3) laboratories and animal housing facilities at APHA, meeting Advisory Committee on Dangerous Pathogens level 3 (ACDP-3) and Specified Animal Pathogens Order level 4 (SAPO-4) requirements. All birds had access to food and water *ad libitum*.

## Acknowledgements

The authors would like to thank Dr. Craig Ross for technical advice and assistance inoculating the pigeons, and colleagues in the Animal Sciences Department at APHA for animal husbandry, clinical assessment and sample collection.

## Author contribution

Conceptualisation: MJS, AB, JJ; formal analysis: CDG, CJW, MJS, JJ; investigation: CDG, CJW, SJ, SR, KR, ST, ALS, AN, DJ, KR, JH, EB; resources: AN, MJS, AB, JJ; writing—original draft: CDG, CJW, DJ, MJS, JJ; writing—review and editing: CDG, AN, MJS, JJ, AB, IHB, AN, ALS. All authors have read and agreed to the final version of the manuscript.

