## Supplementary Figures and Tables for "Pigeons exhibit low susceptibility and poor transmission capacity for H5N1 clade 2.3.4.4b high pathogenicity avian influenza virus"

### Table S1

**Supplementary Table 1.** Summary of observed clinical signs in pigeons and chickens, following initial pigeon inoculation with H5N1-AB.

| | **Total counts of clinical sign occurrence** | | | | | | | | |
|---|---|---|---|---|---|---|---|---|---|
| | **Low dose ($10^2$ $EID_{50}$)** | | | **Medium dose ($10^4$ $EID_{50}$)** | | | **High dose ($10^6$ $EID_{50}$)** | | |
| **Clincial Sign** | **D0 Pigeon** | **R1 Pigeon** | **R1 Chicken** | **D0 Pigeon** | **R1 Pigeon** | **R1 Chicken** | **D0 Pigeon** | **R1 Pigeon** | **R1 Chicken** |
| Changes in huddling | 0 | **1** | 0 | 0 | 0 | 0 | 0 | 0 | 0 |
| Eyes closed | 0 | 0 | 0 | 0 | 0 | 0 | 0 | 0 | 0 |
| Conjunctivitis | 0 | 0 | 0 | 0 | 0 | 0 | 0 | 0 | 0 |
| Changes in body position | 0 | **1** | 0 | 0 | 0 | 0 | **1** | 0 | 0 |
| Oedema | 0 | 0 | 0 | 0 | 0 | 0 | 0 | 0 | 0 |
| Cyanosis of extremities | 0 | 0 | 0 | 0 | 0 | 0 | 0 | 0 | 0 |
| Lethargy | 0 | 0 | 0 | 0 | 0 | 0 | 0 | 0 | 0 |
| Lack of engagement with enrichment | 0 | 0 | 0 | 0 | 0 | 0 | 0 | 0 | 0 |
| Oronasal discharge | 0 | 0 | 0 | 0 | 0 | 0 | 0 | 0 | 0 |
| Diarrhoea | 0 | 0 | 0 | 0 | 0 | 0 | 0 | 0 | 0 |
| Perceived weight reduction | 0 | 0 | 0 | 0 | 0 | 0 | 0 | 0 | 0 |
| Loss of balance | 0 | 0 | 0 | 0 | 0 | 0 | 0 | 0 | 0 |
| Tremors | 0 | 0 | 0 | 0 | 0 | 0 | 0 | 0 | 0 |
| Torticollis | 0 | 0 | 0 | 0 | 0 | 0 | 0 | 0 | 0 |
| Seizure | 0 | 0 | 0 | 0 | 0 | 0 | 0 | 0 | 0 |
| Paralysis/inability to eat or drink | 0 | 0 | 0 | 0 | 0 | 0 | 0 | 0 | 0 |
| **Total** | 0 | **2** | 0 | 0 | 0 | 0 | **1** | 0 | 0 |

D0, directly inoculated pigeons; R1, contact exposed pigeons or chickens.

### Fig. S1

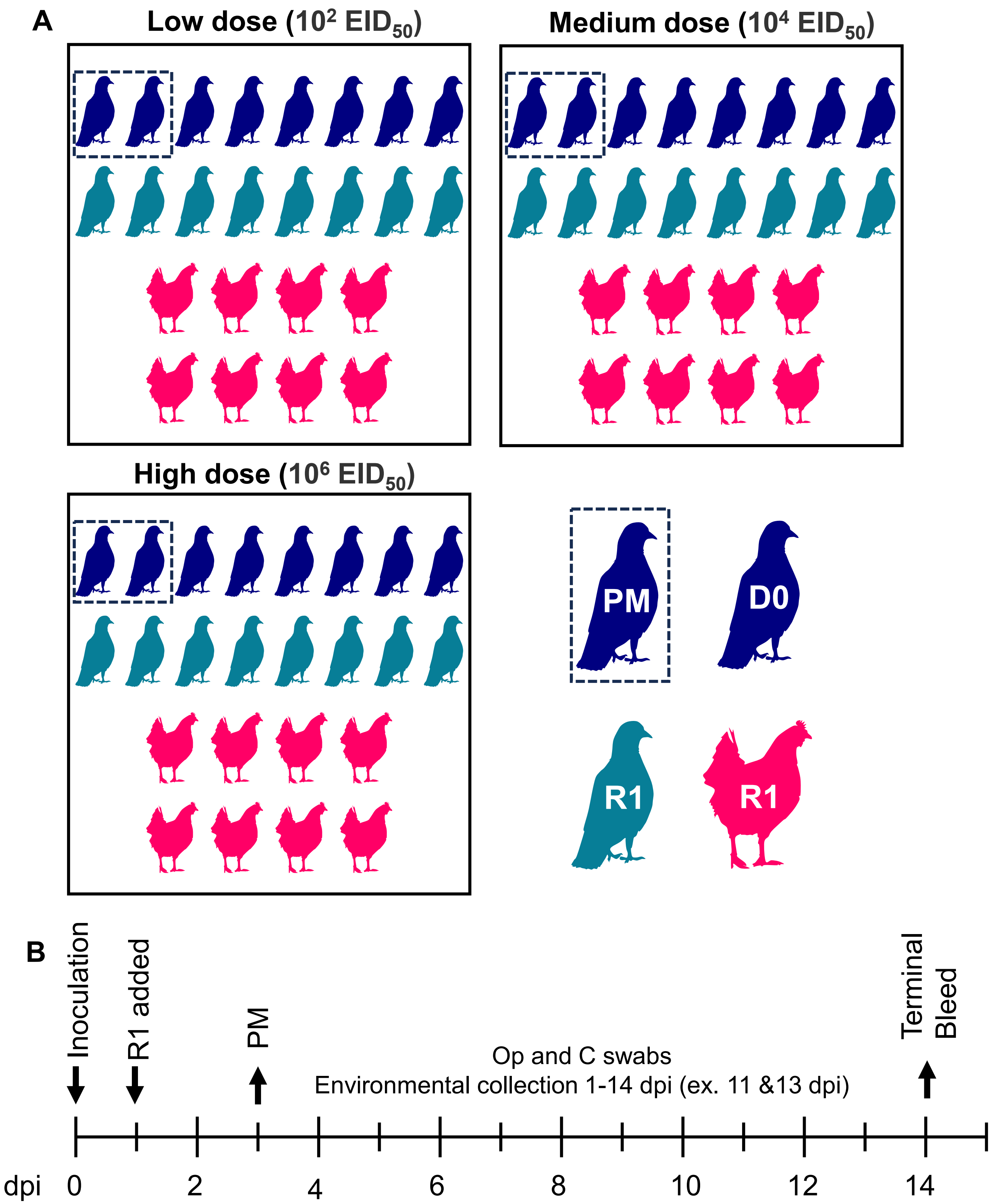


**Fig. S1. Schematic diagram of the pigeon infection and transmission study design.**
**A.** Schematic of the study design. Eight pigeons were directly infected (D0, blue) with low ($10^2$ $EID_{50}$), medium ($10^4$ $EID_{50}$) or high ($10^6$ $EID_{50}$) doses of H5N1-AB. Eight pigeons (teal) and eight chickens (pink) were co-housed (R1) with the D0 pigeons. Two D0 were culled for postmortem (PM) analysis (hashed box). **B.** timeline of the study showing key events with days post infection (dpi) shown.

Fig. S2

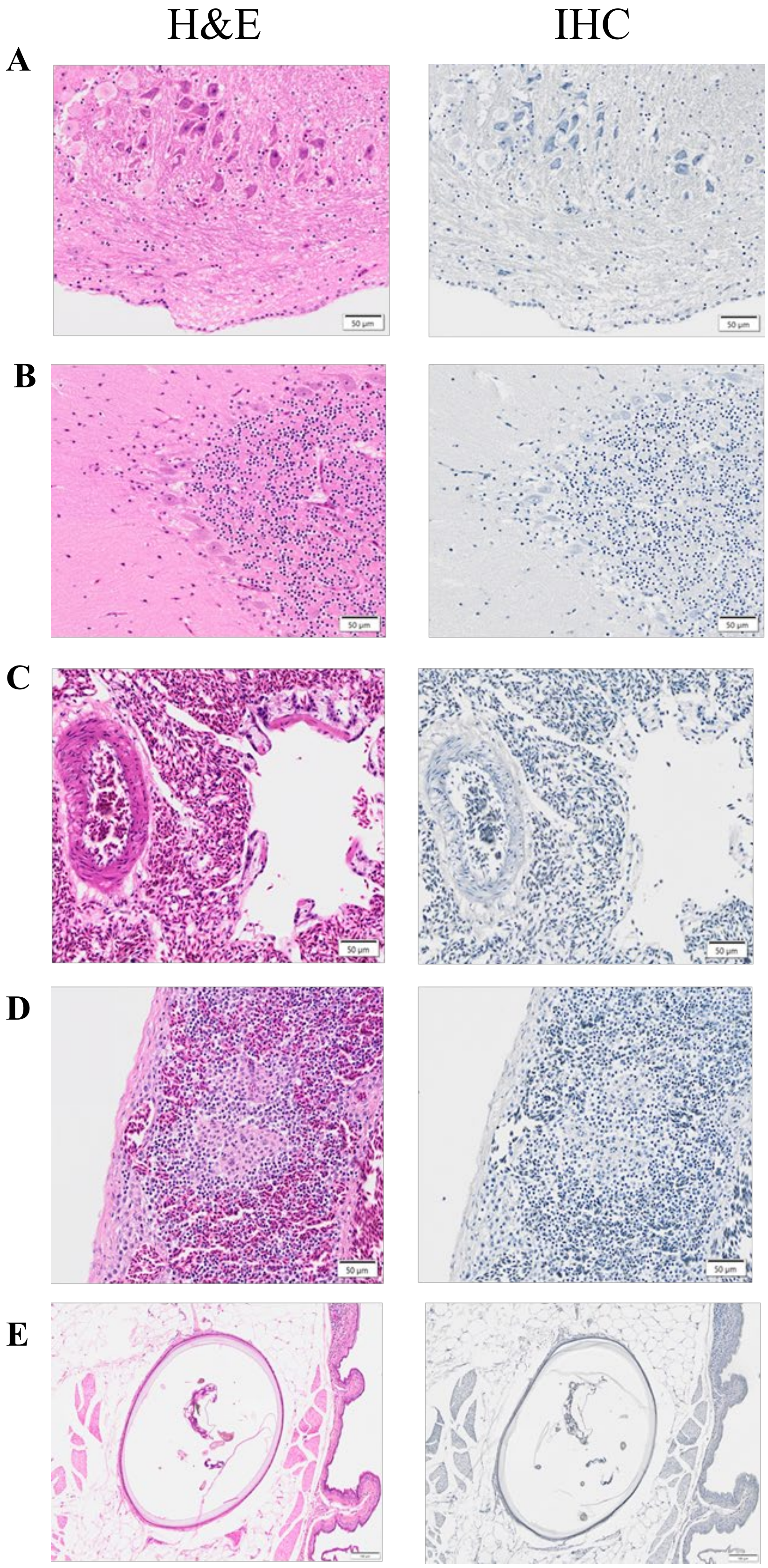


**Fig. S2. Representative tissues sections used for pathological and influenza A viral nucleoprotein (NP) staining, following pigeon infection with H5N1-AB.**

Representative images taken from scanned slides from pigeon #41 (3 dpi) and edited by Olympus Olyvia software. Tissue sections from the same location were stained with haematoxylin and eosin (H&E) (left) or used for virus-specific immunohistochemistry (IHC) (viral nucleoprotein would stain brown but was not detected in any tissue) (right). (**A**) Cerebrum (neurons, oligodendrocyte, microglial cells). (**B**) cerebellum (granule cell layer on right-hand side and molecular layer on the left with Purkinje neurons in-between). (**C**) spleen (red and white pulp). (**D**) lung showing major blood vessel (left-hand side) and parabronchus (right-hand side). (**E**) epidermis, dermis, subcutis and feather pulp.

### Fig. S3

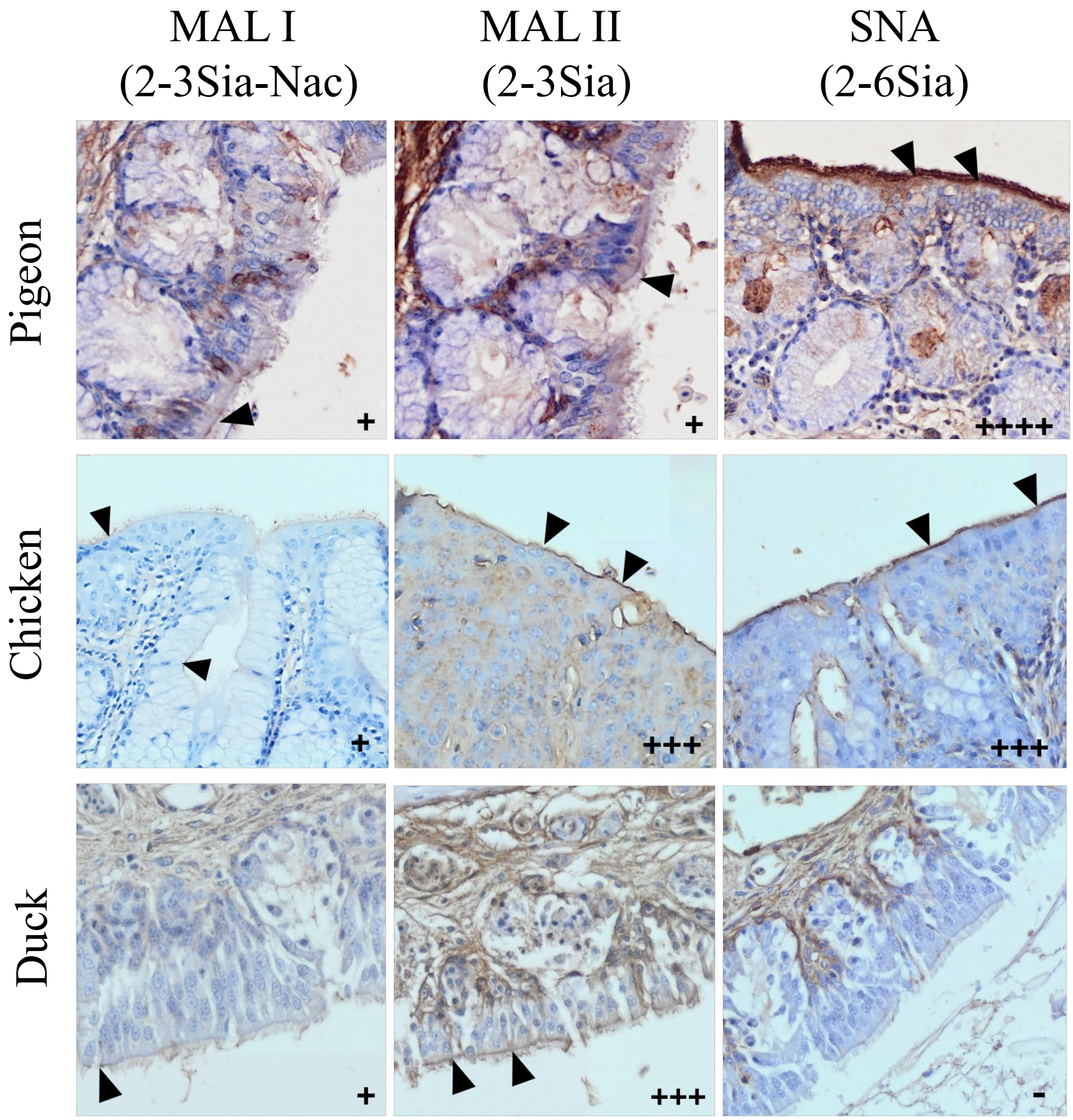


**Fig. S3. Representative images of lectin histochemical labelling in the nasal turbinates of chickens, ducks and pigeons.** Lectin staining demonstrates the distribution and relative abundance (from none (-) to abundant (++++)) of 2-3Sia and 2-6Sia receptors. The presence of receptors is demonstrated on the apical surface of the ciliated epithelial cells as brown line (black arrowheads). Data from these representative images across a range of additional respiratory tissues is summarised in Table 1.
